# DeePaC: Predicting pathogenic potential of novel DNA with reverse-complement neural networks

**DOI:** 10.1101/535286

**Authors:** Jakub M. Bartoszewicz, Anja Seidel, Robert Rentzsch, Bernhard Y. Renard

## Abstract

**Motivation:** We expect novel pathogens to arise due to their fast-paced evolution, and new species to be discovered thanks to advances in DNA sequencing and metagenomics. Moreover, recent developments in synthetic biology raise concerns that some strains of bacteria could be modified for malicious purposes. Traditional approaches to open-view pathogen detection depend on databases of known organisms, which limits their performance on unknown, unrecognized, and unmapped sequences. In contrast, machine learning methods can infer pathogenic phenotypes from single NGS reads, even though the biological context is unavailable.

**Results:** We present DeePaC, a Deep Learning Approach to Pathogenicity Classification. It includes a flexible framework allowing easy evaluation of neural architectures with reverse-complement parameter sharing. We show that convolutional neural networks and LSTMs outperform the state-of-the-art based on both sequence homology and machine learning. Combining a deep learning approach with integrating the predictions for both mates in a read pair results in cutting the error rate almost in half in comparison to the previous state-of-the-art.

**Availability:** The code and the models are available at: https://gitlab.com/rki_bioinformatics/DeePaC

**Supplementary information:** Supplementary data are available at *Bioinformatics* online.

## 1 Introduction

### 1.1 Motivation

Bacterial pathogens evolve quickly and the globalized world enables fast transmission of causative agents over great distances. Virulence genes may be easily exchanged between many bacterial species, leading to emergence of new biological threats. In 2011, for example, a strain of *Escherichia coli* that acquired Shiga-toxin producing genes caused a major outbreak killing 53 people (Frank *et al.*, 2011).

Since DNA sequencing has become the state-of-the art in open-view pathogen detection (Lecuit and Eloit, 2014; Calistri and Palù, 2015), new pipelines and algorithms are needed to efficiently and accurately process the wealth of data resulting from every run. The number of sequences deposited in public databases grows exponentially, and a new challenge is to design computational tools for dealing with big sets of very divergent sequences (Piro *et al.*, 2018). Furthermore, up to a trillion microbial species may be inhabiting the planet (Locey and Lennon, 2016). If this debated estimate (Willis, 2016) is correct, about 99.999% of the microbial biodiversity remains to be discovered. Both yet-unknown and newly-emerging organisms may pose a public health threat. The risks are difficult to anticipate, but quick assessment is mandatory.

Recent developments in the field of synthetic biology have raised concerns about the possibility of creating new biological threats in the lab, either by accident or for malicious purposes. The National Academies of Sciences, Engineering, and Medicine (2018) identify genetic modification of existing bacteria to make them more dangerous as an issue of the highest concern. New methods for sequence-based classification of potential pathogens must be developed to safeguard future biosecurity and biosafety alike (National Research Council, 2010). Approaches based on comparing sequences to a reference database to detect previously known organisms are insufficient. This is especially important for shorter sequences, like next-generation sequencing (NGS) reads or synthetic oligonucleotides. The latter are often not screened before synthesis due to high computational cost and low accuracy of the predictions (Carter and Friedman, 2015; National Academies of Sciences, Engineering, and Medicine, 2018).

Assessing and mitigating risks based on the DNA sequence alone should involve computational methods able to recognize relevant patterns and generate predictions for novel sequences. Therefore, machine learning based approaches are a promising alternative to the traditional sequence analysis tools. For example, deep convolutional networks can be used to predict the lab of origin of the plasmids available in the Addgene repository (Nielsen and Voigt, 2018). This could help track a biological threat back to its source in case of a malicious attack or accidental release. Deneke *et al.* (2017) presented PaPrBaG, a random forest approach for predicting whether an Illumina read originates from a pathogenic or a non-pathogenic bacterium and showed that it generalizes to novel, previously unseen species. They introduce the concept of a *pathogenic potential* to differentiate between predicted probabilities of a given phenotype and true pathogenicity, which can only be realized in the biological context of a full genome and a specific host.

In this work, we present DeePaC, a <u>Dee</u>p Learning Approach to <u>Pa</u>thogenicity <u>C</u>lassification. We focus on the scenario of pathogen detection from next-generation sequencing data. However, the method presented here can in principle be used also for other sequences of similar length, both natural and synthetic.

### 1.2 Computational tools for pathogen detection

#### 1.2.1 Taxonomy-dependent

Read-based pathogen detection methods may be roughly divided in two categories. Taxonomy-dependent approaches directly rely on lists and databases of known pathogens, aiming at assigning sequences to taxonomic categories. Read mappers, for example BWA (Li and Durbin, 2009) and Bowtie2 (Langmead and Salzberg, 2012) fall into this category. Live mapping approaches such as HiLive and HiLive2 (Lindner *et al.*, 2017; Loka *et al.*, 2018) can even map the reads in real time, as the sequencer is running, leading to a drastic reduction in total analysis time.

In general, read mappers specialize in computationally efficient alignment of NGS reads to reference genomes with high specificity. For this reason, they are routinely used for detection of known pathogens, but do not perform well when a sample contains organisms absent from the reference index. Specialized pipelines (Hong *et al.*, 2014; Andrusch *et al.*, 2018) use read mappers in conjunction with additional filtering steps for accurate diagnostics for clinical samples. BLAST (Altschul *et al.*, 1990) offers much more sensitive alignment, appropriate for between-species comparisons, but at the cost of much lower throughput.

Metagenomic profiling tools may also be used as taxonomy-dependent pathogen detection methods. Kraken (Wood and Salzberg, 2014) builds a database of unique 31-mers to assign reads to their corresponding taxonomic units. Ambiguities are resolved by returning the lowest common ancestor. MetaPhlAn2 (Truong *et al.*, 2015) uses around 1 million clade-specific marker genes to detect sequences matching its reference genomes. MicrobeGPS (Lindner and Renard, 2015) identifies potentially unknown organisms present in a metagenomic sample and estimates genomic distances between them and known references. NBC (Rosen *et al.*, 2008, 2011), a naïve Bayes classifier, is a machine learning based method trained to recognize taxa based on their *k*-mer frequency profiles.

#### 1.2.2 Taxonomy-agnostic

Taxonomy-agnostic methods strive to predict phenotypes directly from DNA sequences, without performing any taxonomic assignment. They are not entirely taxonomy-independent, however, as they must be trained on available references. This may lead to bias with regard to the over- and underrepresented taxa in a training set. One goal of the taxonomy-agnostic approaches is to minimize that bias and offer relatively accurate predictions even for novel and divergent sequences. In contrast, the taxonomy-dependent methods are entirely based on the correspondence between a phenotype and a taxonomic classification.

As NBC allows constructing a custom reference database, it may also be used in a taxonomy-agnostic manner. However, it is outperformed by PaPrBaG (Deneke *et al.*, 2017), a random forest approach using a wide range of *k*-mer and peptide-based features. Despite being a read-based method, it can be also used to predict a phenotype from a whole genome. Any long sequence may be fragmented into read-length subsequences. A mean over all the predictions constitutes the final prediction by majority vote. Although this approach is limited to detecting relatively local patterns and cannot use a wider genomic context, it may be useful in practice. Barash *et al.* (2018) used PaPrBaG, PathogenFinder (Cosentino *et al.*, 2013) and their original tool BacPaCS to predict labels for novel genomes recently deposited in the PATRIC database (Wattam *et al.*, 2017).

### 1.3 Deep learning for DNA sequences

Deep learning (LeCun *et al.*, 2015) has been successfully used on genomic data to detect genome accessibility (Kelley *et al.*, 2016) or transcription factor binding sites and disease-associated variants (Alipanahi *et al.*, 2015; Zhou and Troyanskaya, 2015; Zeng *et al.*, 2016; Quang and Xie, 2016; Greenside *et al.*, 2018). Budach and Marsico (2018) implemented convolutional neural networks (CNNs) and long short-term memory networks (LSTMs) in a recently published package, pysster.

Identical predictions for a given sequence of interest and its reverse-complement can be enforced by passing them both through a network, and merging the predictions for each of the strands with a scalar or element-wise operation. Merging may be based for example on the maximum function (Alipanahi *et al.*, 2015; Qin and Feng, 2017) or averaging the results (Quang and Xie, 2017). Resulting networks have a branched architecture resembling Siamese networks.

Shrikumar *et al.* (2017) showed that using standard CNNs leads to inconsistent predictions, and that even data augmentation with reverse-complement sequences cannot mitigate the problem. They developed a solution based reverse-complement parameter sharing between pairs of convolutional filters. A limitation of this method is that it is only directly applicable to convolutional networks. Analogous approaches were independently proposed by several groups (Cohen and Welling, 2016; Kopp and Schulte-Sasse, 2017; Onimaru *et al.*, 2018), and extended with Bayesian dropout (Brown *et al.*, 2018).

## 2 Methods

### 2.1 Data preprocessing

#### 2.1.1 IMG dataset

Similarly to Deneke *et al.* (2017), we accessed the IMG/M database (Chen *et al.*, 2019) on April 17, 2018, and followed the procedure described to identify pathogenic and non-pathogenic bacteria with a human host. Briefly, we filtered the database searching for keywords resulting in unambiguous assignment of a given strain to one of the classes. Previous research successfully used both permanent draft and finished genomes in a similar setting. We decided to include draft genomes in our dataset, too, as their quality is similar to the quality of permanent drafts. For seven species of well-known pathogens (*Campylobacter jejuni*, *Clostridioides difficile*, *Clostridium botulinum*, *Francisella tularensis*, *Listeria monocytogenes*, *Staphylococcus aureus*, and *Staphylococcus epidermidis*), we found between one and two non-pathogenic strains and multiple pathogenic strains; we removed the non-pathogenic strains from further analysis. For *E. coli*, we found multiple strains of both labels. We decided that a study focusing on pathogens should be rather more than less sensitive to this particularly diverse species, especially since some strains may lead to dangerous outbreaks, like the EHEC epidemic in Germany in 2011 (Frank *et al.*, 2011). Therefore, we also removed the non-pathogenic strains of *E. coli* from the final dataset. The procedure yielded a collection of 2,878 strains (2,796 pathogens and 82 non-pathogens) described by data including species names, NCBI Bioproject IDs, and class labels. We then linked the Bioproject IDs with GenBank assembly accession numbers found in the GenBank assembly summary file downloaded on April 17, 2018 as well. We downloaded the assemblies and selected one strain per species at random to avoid skewing the model’s performance towards species with many sequenced strains available.

#### 2.1.2 Species-level validation

This resulted in a list of 446 species (389 pathogens and 57 non-pathogens), with a genome of a single strain representing each species. We assigned 80% of the species to the training set, and 10% to the validation and test sets each, keeping the ratios between pathogens and non-pathogens the same in all sets. Note that the true distribution of pathogenic and non-pathogenic species is unknown. The observed imbalance is just a result of bias towards studying pathogens and does not reflect the reality. Therefore, using an imbalanced test set would only propagate the bias of the source database. Since we are equally interested in accurate predictions for both classes, we use a balanced test set.

Using the Mason read simulator with the Illumina error model (Holtgrewe, 2010) and a custom R script, we simulated 10 million single-end Illumina reads per class from the training set and 1.25 million reads per class from the validation set, thus balancing the number of data points in both classes. We simulated 1.25 million paired reads per class from the test set, allowing us to evaluate the classification performance in both the single-end and the paired-end setting. This amounts to 80%, 10%, and 10% of all reads assigned to each of the sets, respectively. Read length was set to 250 bases in all cases, representing what is routinely available using an Illumina MiSeq device; for read pairs we used a mean fragment length of 600 with a standard deviation of 60. We also tested whether using an imbalanced training set with class weighting could improve the results, but the corresponding networks were outperformed by those trained on a balanced set and were not selected as our final models. For a more detailed description of the class-weighted training setting, see Supplementary Information.

#### 2.1.3 Read-level validation

Note that in our primary dataset we placed separate species in training and validation sets to explicitly force our classifiers to predict pathogenic potentials of reads originating from *novel* species. A simpler approach would comprise simulating the reads first, and then assigning them to the training and validation sets. However, this would lead to solving a similar, but different biological problem (classifying reads originating from *known* organisms). To explicitly test those assumptions, we generated a version of the training and validation set where reads from the same species occur in both training and validation sets. In this setting, the test set remained the same as above, so it would be possible to compare the effects of read-level and species-level validation.

#### 2.1.4 Temporal hold-out data

We accessed the IMG/M database again on January 17, 2019 and preprocessed it as described above. We identified three new species passing the filters applied. All of them were pathogens belonging to the *Pantoea* genus, which is absent from our original dataset. We downloaded the genomes and simulated paired reads as described above. To keep the mean coverage at a similar level to the coverage of the original test set, we simulated 100,000 reads in total. While this supporting dataset is small, it is useful as a case study in conjuction with the results of evaluation on the more diverse main dataset.

#### 2.1.5 Real data case study

To test the performance of our classifier on data from a real sequencing run, we searched the SRA database for sequencing runs performed for a well-known pathogenic species *Staphylococcus aureus*. This species was not present in the training set (it had been randomly assigned to the validation set). We considered runs performed on the Illumina platform with paired-end reads of length 250. Note that in case of real sequencing reads, the true read length may vary, and some of the reads were significantly shorter. We accessed the archive SRX4814864, originating from a lesion swab from an Italian paediatric hospital (Manara *et al.*, 2018). We downloaded the data from the corresponding run SRR7983698 as unclipped FASTA files.

#### 2.1.6 BacPaCS dataset

Finally, we downloaded all the data used by Barash *et al.* (2018) in the assessment of their BacPaCS method. We accessed the PATRIC database (Wattam *et al.*, 2017) by the IDs provided by Barash *et al.* (2018), copying their training and test sets for direct comparability. The original training set consists of 17,811 genomes of pathogens and 3,274 genomes of non-pathogens, while the test set contains 60 pathogens and 40 non-pathogens. Importantly, those genomes do not represent unique species and the number of strains per species varies greatly. However, following Barash *et al.* (2018), we treated each strain as a separate entity and used all of them in the analysis. We randomly reassigned 10% of the original training genomes to the validation set, so our BacPaCS training set comprised of 90% of the original. The test set was left unchanged. The read simulation was then performed exactly the same as for the species-level validation.

### 2.2 Reverse-complement networks

#### 2.2.1 DNA encoding and reverse-complementarity

We use distributed orthographic representations of DNA sequences, a method based on one-hot encoding of every nucleotide in the sequence. Namely, a sequence is converted into a 2D binary tensor, where one dimension represents the position of a given nucleotide in the sequence, and the other represents the one-letter nucleotide code as a one-hot encoded vector. For example, adenine is represented by the vector (1,0,0,0) and thymine by (0,0,0,1). Unknown nucleotides may be encoded as (0,0,0,0). Note that reversing the sequence tensor along both axes results in the reverse-complement.

We design networks with two separate branches processing each of the input orientations respectively. Each of the branches consists of identical layers and all the parameters are shared between each pair of layers. We propose two variants of this architecture. In the *full* RC-networks, input to the deeper layers consists of concatenated outputs of both the forward and reverse-complement versions of the previous layer. The output of the RC layer is flipped before concatenating, so that the channels *i* and *n −* 1 *− i* in the final tensor correspond to the same feature on opposing strands. Note that in this case, a *full* RC-CNN is equivalent to the RC-CNNs proposed by Shrikumar *et al.* (2017). In the second variant, dubbed a *Siamese* RC-architecture, each of the branches functions separately before the merging layer. That means that the input to a deeper layer is just the output of the previous layer in a branch. We tested both variants using CNNs, bidirectional LSTMs and hybrid networks with both convolutional and LSTM layers, but they are in principle compatible with any other neural architecture. We also considered three methods of strain representation merging. Although the differences were small, a summation of forward and reverse-complement feature vectors yielded the best performance overall. For a more detailed description of the merging methods used, see Supplementary Information.

#### 2.2.2 Species-level and paired reads predictions

One of the major challenges of pathogenic potential prediction from single reads is the lack of biological context. However, if all the reads in a sample originate from the exactly same organism, we can predict the pathogenic potential of that organism by a majority vote. In the context of probabilistic estimates of the class label (returned by both PaPrBaG and our neural networks), we can implement that as a simple mean over predictions for all the individual reads. For BLAST, we can just assign the label predicted for the majority of reads.

Building upon this idea, we can boost read-based performance if we consider read pairs, assumed to originate from the same organism even in metagenomic samples. To this end, we average predictions for the corresponding pairs in our test set. The classifiers may still predict pathogenic potentials for isolated sequences if so desired. We can integrate binary predictions (e.g. returned by BLAST), taking into account the missing and conflicting predictions for some of the reads. We treat missing predictions as undefined values and implement the *accept anything* operator of ternary logic. It returns a positive label if and only if one of the input values is positive, and the other is not negative. Conversely, it returns a negative label if and only if one of the input values is negative, and the other is not positive. The result is undefined when both inputs are undefined, or in case of conflicting input values.

#### 2.2.3 Hyperparameter tuning

We used Keras 2.2.4 and TensorFlow 1.12. We tested a total of 243 different architectures. All were initialized with He weight initialization (He *et al.*, 2015) and trained with the Adam optimizer (Kingma and Ba, 2014). We used dropout regularization (Srivastava *et al.*, 2014), including input dropout (interpreted as setting a random fraction of nucleotides to Ns). We also applied batch normalization (Ioffe and Szegedy, 2015) to some of the RC-CNNs and tested the effect of adding an L_2_ regularization term in RC-LSTMs. In addition, we trained traditional CNNs and LSTMs without RC parameter sharing, equivalent to the networks that can be implemented with the pysster package (Budach and Marsico, 2018). Finally, we selected the best CNN and the best LSTM model, and prepared a simple ensemble classifier by averaging the predictions of those two models. For a more detailed description of the tuning process, see Supplementary Information.

#### 2.2.4 Read-level validation

Generalization from one set of reads to another set of reads should be relatively easy when both sets originate from the same species. However, we expected that even very high read-level validation accuracy would not translate into high test accuracy (see section 2.1.3). After the primary tuning procedure described in the previous sections, we selected the architecture obtaining the highest *training* accuracy, as we assumed little overfitting would be seen at the validation stage in this setting. We trained the network using read-level validation and evaluated it on our primary test set containing reads from species absent in both validation and training sets.

### 2.3 Benchmarking

#### 2.3.1 PaPrBaG

To benchmark our method against the state-of-the art in pathogenic potential prediction, we trained PaPrBaG random forests on our training set, and evaluated them on our test set. We used two different feature settings for PaPrBaG. The original authors extracted more than 900 features from each of the reads, including *k*-mer frequencies and a selection of amino-acid and peptide features inferred by a “least STOPs” heuristic. However, they show that the translation-based features contribute relatively little to the final classification decision. Therefore, we decided to use both the original feature set, and a reduced set comprising the DNA features only.

#### 2.3.2 BLAST

BLAST was shown (Deneke *et al.*, 2017) to achieve the best read-by-read performance among alternatives to machine learning approaches for predicting pathogenic potential. It outperformed both the mapping-based Bowtie2 and Pathoscope2, which builds on the former, as well as two different variants of the *k*-mer based Kraken (Langmead and Salzberg, 2012; Hong *et al.*, 2014; Wood and Salzberg, 2014). All three failed to classify most of the reads. Furthermore, BLAST achieved better classification results than a naïve Bayes classifier (NBC) with a word length of 15 (Rosen *et al.*, 2011).

For each test read, we performed a search against the database containing all the training genomes and take a label of the best hit for each of the test reads as a predicted label for that read. We also tested a variant of this approach using all the available strains of the training species to build the database. In both cases, we use the discontiguous megablast task (*-task dc-megaBLAST*), a cutoff E-value of 10 and the default parameters.

#### 2.3.3 BacPaCS

Finally, we compared our approach to BacPaCS, Pathogenfinder, and PaPrBaG using the original BacPaCS test dataset. Without any further tuning, we selected one CNN and one LSTM architecture which worked best on our data. We retrained them both using the BacPaCS training data. Since the BacPaCS dataset treats strains, not species, as the primary entities (and is used by the authors to predict labels for new strains of species *present* in the training set), the classifier designed by Barash *et al.* (2018) can be treated as specialized in predicting pathogenic potentials for *known* species. Therefore, we assumed that the architecture used for read-based validation would yield high performance, and retrained it on the BacPaCS data as well. We evaluated the networks in a single-species sample setting and compared the results to the originally presented performance metrics for BacPaCS and Pathogenfinder.

Assessing PaPrBaG, Barash *et al.* (2018) used the random forests trained on the original PaPrBaG dataset. This may lead to inaccurate estimates of the classification error, as the labels they mined from the PATRIC database differ from the labels that the original PaPrBaG forests used. This problem is exacerbated by the imbalance between the number of strains per species in the BacPaCS test dataset. For example, strains of *Acinetobacter baumannii* alone constitute 30% of the non-pathogens in the set. *Fusobacterium periodonticum* amounts to another 10%. Importantly, both of those two species were treated as pathogens in the original PaPrBaG dataset, but are assigned a non-pathogenic label based on the PATRIC metadata. One should therefore expect that the original PaPrBaG forests will predict wrong labels for a significant fraction of the test set if not retrained using the labels extracted by Barash *et al.* (2018) for their training set. Therefore, we retrained PaPrBaG on the original BacPaCS dataset for an accurate comparison to our networks.

## 3 Results

### 3.1 Pathogenic potential prediction from NGS reads

#### 3.1.1 Single reads

We present the results of evaluation on single NGS reads in Table 1. The values given are calculated over the complete test set, treating each read as a separate entity. Precisely, we can calculate the performance measures for the sets of “left” and “right” mates separately and then compute the mean for any given measure. Results obtained for the “left” and “right” half of the set did not differ from those presented in Table 1 by more than 0.001.

**Table 1.**
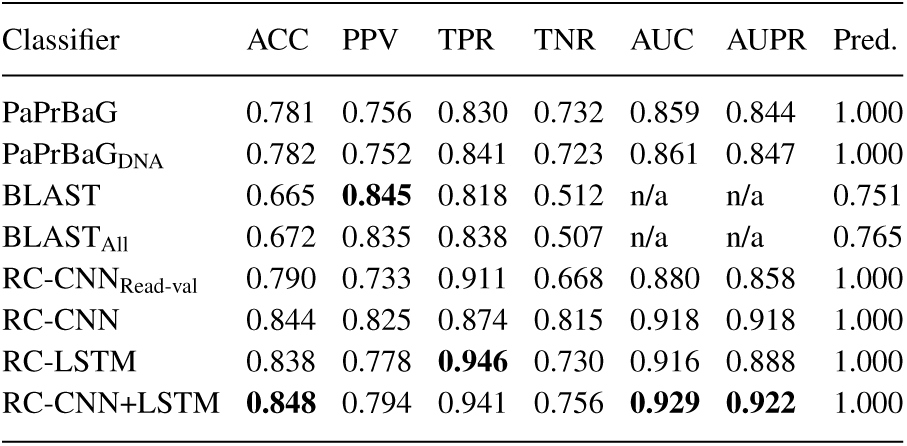
Classification performance on single reads. PaPrBaG_DNA_ is a variant of PaPrBaG using DNA-based features only. By default, we use the training set to build the BLAST reference database. BLAST_All_ uses all available strains of the training species. RC-CNN_Read-val_ was trained using read-level validation. RC-CNN+LSTM is the average of RC-CNN and RC-LSTM predictions.

As previously shown (Deneke *et al.*, 2017), BLAST fails to classify some of the reads at all. To compare its performance to the machine learning approaches, we define accuracy as the ratio of correct predictions to the number of all data points in a set. Therefore, missing predictions are counted as false positives or false negatives depending on the ground truth label. The fraction of classified reads is presented in the last column.

The neural networks clearly outperform both BLAST and PaPrBaG in terms of prediction accuracy. The selected RC-CNN model consists of 2 convolutional and 2 dense layers with 512 and 256 units respectively; it was trained with an input dropout rate of 0.25 and without batch-normalization. The RC-LSTM has one layer of 384 units, and was trained with input dropout of 0.2. The traditional deep learning architectures performed worse than their reverse-complement counterparts (data not shown). However, the differences in validation accuracy were below 0.01. Figure S1 compares the prediction distributions of standard and RC-networks, showing the importance of employing the RC models, especially for LSTMs.

The RC-CNN model achieves the highest positive predictive value (precision), and the RC-LSTM is the most sensitive. The RC-CNN+LSTM ensemble model aims at a trade-off between those two performance measures. Integrating signals contained in both mates significantly boosts performance of all the evaluated methods (Table 2). In this case, the RC-CNN achieves the highest accuracy. Compared to the previous state-of-the-art (PaPrBaG with original settings, single reads only), it cuts the error rate almost in half (21.9% vs. 11.3%).

**Table 2.**
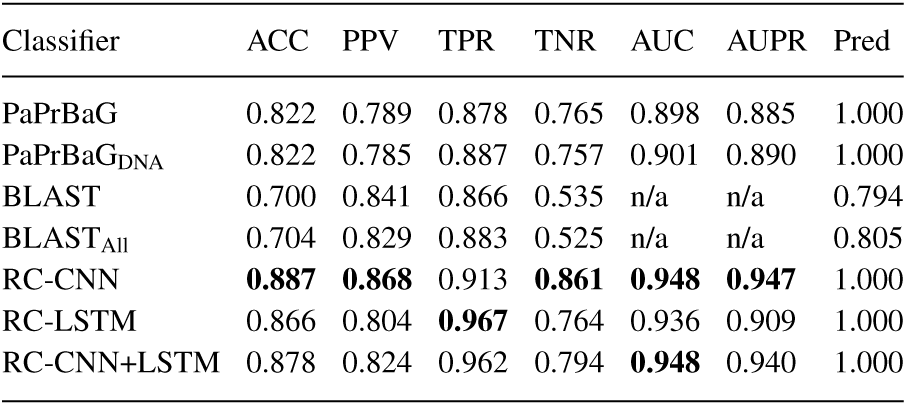
Classification performance on read pairs

A *full* RC-CNN with 2 convolutional and 2 dense layers of 512 and 256 units, batch normalization and no input dropout achieved the highest *training* accuracy and was used for read-level validation. As expected, it performs much worse, even though it achieved an impressive validation accuracy of 0.956. Somewhat surprisingly, it still outperforms both PaPrBaG and BLAST. It is important to note that for all the other methods, the validation accuracy was much lower than test accuracy. It ranged from 0.574 (BLAST) to 0.784 (RC-LSTM) and 0.787 (RC-CNN+LSTM). We hypothesize that this is because the species randomly assigned to the training set are more closely related to the species in the test set than to those in the validation set. We comment on this problem further in the Discussion.

We note that PaPrBaG_DNA_ is faster than the original PaPrBaG at no performance cost. GPU-accelerated networks are even faster and can predict up to 1817 reads/s (RC-LSTM) and 4010 reads/s (RC-CNN; see Supplementary Information for additional details). A million-read sample may be processed in just over 4 minutes even on a consumer-level desktop computer.

#### 3.1.2 Single-species samples

The evaluation scenario presented in Table 3 models sequencing a pure, single-species isolate, when it can be assumed that all reads originate from the same genome. After averaging the pathogenic potentials over whole genomes, BLAST, PaPrBaG, and RC-CNN performed equally well, with just one false positive and one false negative. The RC-LSTM and RC-CNN+LSTM ensemble models predict one more false positive, which is reflected in their drastically lower specificity. This is in line with the previous results suggesting that those models are more sensitive, even though they do not manage to limit the number of false negatives to zero. Note that the set contained 39 pathogen and only 6 non-pathogen genomes. A bigger and more balanced dataset would allow more reliable error estimates, but collecting more data that could be used in a rigorous, automated data preprocessing workflow poses a non-trivial challenge. It is nevertheless important to evaluate prediction accuracy also on single-organism samples, as this is one of the possible applications of the proposed method (Deneke *et al.*, 2017; Barash *et al.*, 2018). Validation balanced accuracy ranged from 0.808 (All-strains BLAST) to 0.891 (BLAST, RC-LSTM). Interestingly, the RC-LSTM and RC-CNN+LSTM ensemble models achieved higher balanced accuracy on the validation set than on the test set. This may reflect the different evolutionary distances between the members of each of the sets, as discussed for the single-read predictions.

**Table 3.** Classification performance on single-species samples. Balanced accuracy (BACC) is the mean of sensitivity (TPR) and specificity (TNR).

| Classifier | ACC | BACC | PPV | TPR | TNR | AUC | AUPR |
| --- | --- | --- | --- | --- | --- | --- | --- |
| PaPrBaG | **0.956** | **0.904** | **0.974** | 0.974 | **0.833** | 0.932 | **0.988** |
| PaPrBaG<sub>DNA</sub> | **0.956** | **0.904** | **0.974** | 0.974 | **0.833** | 0.932 | **0.988** |
| BLAST | **0.956** | **0.904** | **0.974** | 0.974 | **0.833** | n/a | n/a |
| BLAST<sub>All</sub> | **0.956** | **0.904** | **0.974** | 0.974 | **0.833** | n/a | n/a |
| RC-CNN | **0.956** | **0.904** | **0.974** | 0.974 | **0.833** | **0.940** | **0.989** |
| RC-LSTM | 0.933 | 0.821 | 0.950 | 0.974 | 0.667 | 0.897 | 0.978 |
| RC-CNN+LSTM | 0.933 | 0.821 | 0.950 | 0.974 | 0.667 | 0.923 | 0.985 |

### 3.2 Temporal hold-out

For the temporal hold-out test, we predicted labels for reads originating from *Pantoea brenneri*, *Pantoea septica*, and *Pantoea conspicua*, which were labeled pathogenic based on the metadata extracted from the IMG/M database. They were not available when we prepared our training, validation, and primary test sets. Species-wide, the labels were predicted correctly for all three genomes by both BLAST and the reverse-complement networks (Table 4). The RC-LSTM and the RC-CNN+LSTM ensemble model performed best, but the RC-CNN outperformed BLAST as well. PaPrBaG was not used here, as it consistently underperformed in comparison to the deep learning approaches in the previous tests. As mentioned in section *Data preprocessing*, this dataset is too small to yield reliable error estimates and the temporal hold-out should only be interpreted as a case study supporting the other results. The *Pantoea* genus was absent from the training database when the training set was compiled; accurate predictions for a novel genus (while trained at the species level) suggest successful generalization.

**Table 4.** Temporal hold-out, read pairs

| Classifier | ACC | PPV | TPR | Pred. |
| --- | --- | --- | --- | --- |
| BLAST | 0.881 | 1.000 | 0.881 | 0.883 |
| BLAST<sub>All</sub> | 0.895 | 1.000 | 0.895 | 0.896 |
| RC-CNN | 0.984 | 1.000 | 0.984 | 1.000 |
| RC-LSTM | **0.992** | 1.000 | **0.992** | 1.000 |
| RC-CNN+LSTM | **0.992** | 1.000 | **0.992** | 1.000 |

### 3.3 Real-data results

One of the defining characteristics of real sequencing runs is that the reads generated are not all of the same length. Since the neural networks require a constant input length, we pad the sequences shorter than 250nt with zeroes (interpreted as Ns). If any sequence is longer than this threshold, it is trimmed. This may be a problem for the CNNs with average pooling, as multiple zero-entries significantly lower the activations after a pooling layer. Therefore, we expected those architectures to achieve lower accuracy than the RC-LSTMs on a real dataset. Note that this problem does not apply to RC-CNNs with max pooling. However, those performed worse than their average-pooling counterparts in the tuning and validation step. As they essentially work as motif detectors, we suspect that they are more prone to overfitting. We present the results in Table 5.

**Table 5.** Performance on real data, read pairs

| Classifier | ACC | PPV | TPR | Pred. |
| --- | --- | --- | --- | --- |
| BLAST | 0.746 | 1.000 | 0.746 | 0.750 |
| BLAST<sub>All</sub> | 0.787 | 1.000 | 0.787 | 0.790 |
| RC-CNN | 0.900 | 1.000 | 0.900 | 1.000 |
| RC-LSTM | **0.984** | 1.000 | **0.984** | 1.000 |
| RC-CNN+LSTM | 0.982 | 1.000 | 0.982 | 1.000 |

Although the RC-CNN suffers from lower accuracy compared to the RC-LSTM and RC-CNN+LSTM ensemble models, it still outperforms BLAST by a large margin. Based on the prediction speed we can name the RC-CNN our *rapid* model, and the RC-LSTM our *sensitive* model. Their predictions may be aggregated with the RC-CNN+LSTM ensemble model, with almost no additional computation, for a boost in sensitivity at a smaller precision cost.

### 3.4 BacPaCS dataset results

We present the results of evaluation performed on the BacPaCS dataset in Table 6. The RC-networks outperform all of the other methods in terms of balanced accuracy. PaPrBaG’s specificity is much higher after retraining (see section 2.3.3). This is also reflected in balanced accuracy, which in this case is even higher than BacPaCS’s. The RC-CNN architecture selected based on our primary dataset turns out to be the most specific architecture when trained on the BacPaCS dataset. As expected, the best overall performance is achieved by an RC-CNN with batch normalization and without input dropout, as observed for read-level validation (see section 2.2.4).

**Table 6.** Performance on the BacPaCS dataset. BACC is the mean of TPR and TNR.

| Classifier | TPR | TNR | BACC |
| --- | --- | --- | --- |
| BacPaCS | 0.92 | 0.48 | 0.70 |
| Pathogenfinder | 0.63 | 0.28 | 0.46 |
| PaPrBaG (Barash *et al.*) | **1.00** | 0.05 | 0.53 |
| PaPrBaG (retrained) | 0.72 | 0.83 | 0.77 |
| RC-CNN | 0.70 | **0.93** | 0.81 |
| RC-LSTM | 0.75 | 0.85 | 0.80 |
| RC-CNN+LSTM | 0.72 | 0.90 | 0.81 |
| RC-CNN+BN | 0.78 | 0.90 | **0.84** |

## 4 Discussion

### 4.1 Evolutionary distances and pathogenicity

Given the difficulty of the task, the reverse-complement networks offer impressive performance. They are able to predict a high-level, abstract phenotype (namely, pathogenic potential) from isolated reads, without any additional biological context. They also outperformed the traditional deep learning architectures in the tuning and validation step. However, all of the evaluated tools show a noticeable gap between validation and test accuracy. We assume that this is a result of the species-level division of the data into the training, validation and test sets – it must be more difficult to predict correct labels for the validation data than for the test data. We hypothesize that the species assigned to the test set must be more similar (in terms of evolutionary distance) to the training species than the validation species. This discrepancy would be therefore a result of treating all the species as separate, independent entities, as if the sequence similarities between individual genomes (and the corresponding reads) were uniform. This is not biologically correct; closely related species have similar genomes, and horizontal transfer of genetic elements (including virulence factors) introduces local similarities.

Pathogenic and non-pathogenic strains may belong to the same species. However, the extracted labels are actually consistent among individual strains for a vast majority of the species in the IMG/M database. Seven well-known pathogens had between one and two non-pathogenic strains that were ignored in the further analysis. We select one strain per species to avoid skewing the performance towards species with more strains available. As expected, we found multiple *E. coli* strains of both classes. We decided to only consider pathogenic ones, so we assume our models to classify any *E. coli* sample as exhibiting pathogenic potential. While this may be seen as a major limitation of the proposed method, it only affects a single species in the dataset. It also reflects the great phenotypic diversity of *E. coli*. Importantly, this problem does not apply to the models trained on the BacPaCS dataset, which includes multiple independently labelled strains per species. DeePaC performs very well also in this setting.

We resort to the oversimplification of treating species as independent since balancing the read sets to account for sequence similarity is not trivial. The same approach was used by PaPrBaG (Deneke *et al.*, 2017). In turn, BacPaCS (Barash *et al.*, 2018) uses individual strains as primary biological entities, which leads to similar problems. Nevertheless, both approaches do achieve satisfactory results. In fact, Deneke *et al.* (2017) have shown that PaPrBaG outperforms taxonomy-dependent methods (BLAST, Kraken, Bowtie2 and Pathoscope2) in two challenging cases: when species belonging to the same genus have different phenotypes, and when a genus is completely absent from the training set (as in our temporal hold-out). This suggests that read-based, taxonomy-agnostic methods like DeePaC and PaPrBaG are actually more robust to the oversimplified assumption of uniform evolutionary distances between species. A future outlook could include investigating in detail how sequence similarities and differences in labels within each taxonomic rank influence the predictions, as well as which parts of a given genome lead to the most confident predictions.

More exact estimates of the classification error could be obtained using nested cross-validation, but this was not computationally feasible – a single training epoch may take up to 6 hours on a state-of-the-art GPU for some of the most demanding architectures. Therefore, we assume that the accuracy of BLAST, a sensitive alignment-based approach, reflects sequence similarities (and presumed evolutionary distances). This means that a method more accurate than BLAST generalizes well. In addition, we test the classifiers on independent data. Our approach consistently outperforms both BLAST and PaPrBaG. It also fares better that BacPaCS, Pathogenfinder, and PaPrBaG when tested on the original BacPaCS data.

### 4.2 Predictions from single reads

The results above may be counter-intuitive. Virulence factors are often encoded on pathogenicity islands or mobile genetic elements. While the genome assemblies used in this study often include plasmids (which can hence be detected by the models), it is impossible to guarantee that none was missed. Moreover, the same organism may cause a disease in one person and not in the other – for those very reasons we can only predict pathogenic potentials, not a pathogenic phenotype directly. All this context is unavailable in an NGS read. It is not even possible to reliably detect open reading frames; comparing PaPrBaG to PaPrBaG_DNA_ proves that attempting to infer putative peptide sequences only introduces noise. Nevertheless, the evaluation clearly shows that the read-based approaches do work. Surprisingly, DeePaC outperforms BacPaCS, which predicts pathogenic potentials from whole proteomes, treating the most relevant genes as input features (Barash *et al.*, 2018). Apparently, predicting pathogenic potentials based on isolated reads and aggregating the predictions with a simple mean results in accurate phenotype predictions – also when multiple strains of each species are considered.

This could be related to the texture-bias of CNNs. Brendel and Bethge (2019) have recently shown that splitting an image into receptive fields as small as 17 × 17 px, and subsequent averaging of the class predictions for each of the patches results in high accuracy predictions on the ImageNet dataset. A similar effect may be seen in the pathogenic potential prediction task. A genome is split into short reads, but the local sequence patterns are sufficient to predict a phenotypic label even though establishing a mechanistic link between a read and the phenotype is improbable. Averaging the results leads to confident predictions for whole genomes in single-organism samples.

### 4.3 The definition of a pathogen

It is important to define human pathogens in a reliable and consistent way. However, it is also by no means a trivial task. We do not differentiate between the level of danger individual organisms may pose – dangerous, biosafety level 3 select-agents are put in the same category as opportunistic pathogens causing relatively mild and easily-treated infections. Classifying the risk posed by unrecognized DNA at a higher resolution is an interesting avenue for further research; it is however likely to be challenging due to a comparatively small number of very dangerous bacterial species. We recognize that what constitutes a pathogen is an open problem and different definitions may be interesting for different purposes. For processing big datasets, the labels must be extracted automatically using handcrafted rules compatible with a particular source database. It is therefore immensely important to be aware of the underlying assumptions, as they will be reflected in the final classification decisions of a trained model. We note that comparing models trained using incompatible labels may result in unreliable error estimates, especially if organisms for which the labels differ constitute a substantial fraction of the test set. We recommend sharing the metadata describing the training organisms along with the trained models. For this study, they are available at https://gitlab.com/rki_bioinformatics/DeePaC.

### 4.4 A flexible framework for RC-constrained classification

DeePaC is easily extensible and may be used as a generalized framework for building reverse-complement neural architectures beyond the applications described here. It is not constrained to the read length used here or the pathogenic potential prediction task; any label or value may be used as a prediction target. For predicting a categorical phenotype directly from NGS reads, we suggest that the target label should be a relatively abstract, general feature. Phenotypes dependent on single genes or small sets of genes will most probably be very difficult to predict from whole-genome sequencing reads. The variant of the RC configuration (*full*, *Siamese* or *none*) may be easily switched, so the framework can be also applied to tasks where the RC-constraint is not important. RNA sequences can be analysed, provided that they are converted to binary tensors using the appropriate alphabet.

## 5 Conclusions

We show that the RC-networks outperform the previous state-of-the-art on both simulated and real sequencing data, accurately predicting pathogenic potentials from isolated NGS reads. The trained models can be used to predict pathogenic potentials for unknown, unrecognized and novel (e.g. synthetic) DNA sequences with a simple script, also if they do not originate from an Illumina sequencing run. It is also possible to filter a read set based on a user-defined threshold to select only the reads for which a network yields confident predictions. This may be used as a crude pre-filter, where the reads with the highest pathogenic potential may be further investigated in downstream analysis. The code, the models trained and example configuration files are available at https://gitlab.com/rki_bioinformatics/DeePaC.

## Acknowledgements

We are grateful to Stefan Budach and Annalisa Marsico (Max Planck Institute for Molecular Genetics) for their insights which helped us design the first prototype CNN. We also thank Carlus Deneke (German Federal Institute for Risk Assessment) for sharing his expertise regarding PaPrBaG. We express our gratitude to Melania Nowicka (Max Planck Institute for Molecular Genetics), Tobias Loka and Vitor C. Piro (Robert Koch Institute) for many fruitful discussions and comments.

