## Supplementary Information for "DeePaC: Predicting pathogenic potential of novel DNA with reverse-complement neural networks"

Bartoszewicz *et al.*

### 1 Representation merging

We note that forward and reverse-complement representations can be merged by any element-wise operation on the 1D output tensors. Similarly to Shrikumar *et al.* (2017), we place the dense layers (and the output layer) after representation merging. Apart from summation, we consider two alternative merging functions (note that averaging and adding the representation vectors are essentially equivalent). The *max* function implements the Gödel $t$-conorm, corresponding to the OR operation in Gödel fuzzy logic. Even though the activations are not restricted to the interval [0,1], high output values can be interpreted as finding a motif on either of the two strands. The Hadamard product is the product $t$-norm corresponding to the AND operation in the product fuzzy logic. Here, high values may be understood as finding a motif on both strands at the same time.

### 2 Training and tuning

#### 2.1 Class weighting

While the data preprocessing procedure results in a balanced training dataset, the mean coverage of pathogen and non-pathogen genomes is drastically different. To explore an alternative way of solving the class imbalance problem, we also simulated an imbalanced training set, where the total of 20 million training reads was simulated with equal mean coverage from all the training genomes, regardless of their labels. This dataset was then used to train ten networks using a class-weighted loss function. PaPrBaG, constituting the state-of-the art in machine learning based pathogenicity prediction, does not support error weighting. For BLAST, a method based on sequence homology, this distinction does not apply at all, as its reference database is constructed over whole genomes. However, this does not influence our final results; the class-weighted networks were outperformed by those trained on a balanced set and were not selected as our final models.

### 2.2 Hardware

Each of the networks was trained on between one and four GPUs (GTX 980, GTX 1080 Ti, or RTX 2080 Ti) depending on hardware availability at a given time point. CUDA 9.0 was used on machines equipped with the GTX-line cards, but the RTX 2080 Ti card required CUDA version 10.0.

### 2.3 Hyperparameter tuning: RC-LSTM

All networks were trained with the Adam optimizer (Kingma and Ba, 2014) using the default parameters and a batch size of 512. We finished the training after a maximum of 15 epochs or earlier if the validation accuracy did not improve for 10 consecutive epochs. We used dropout regularization (Srivastava *et al.*, 2014) with a dropout rate of 0.5 after all recurrent and dense layers, and input dropout in most of the architectures. For the bidirectional RC-LSTMs, we tested the following parameter combinations for both the *full* and the *Siamese* variant:

- sum merge, 1 layer, 128-512 units, input dropout rate: 0, 0.1-0.3
- max merge, 1 layer, 384 units, input dropout rate: 0.1-0.3
- product merge, 1 layer, 384 units, input dropout rate: 0.1-0.3
- sum merge, 2 layers, 384 units, input dropout rate: 0.1-0.3

The input dropout rate was varied with step of 0.05 unless stated otherwise, and the number of units was adjusted with a step of 128. In addition, we trained two traditional LSTMs without RC parameter sharing, with 256-384 units and an input dropout of 0.2.

### 2.4 Hyperparameter tuning: RC-CNN

For the RC-CNN architectures, we used 64-512 units in the convolutional layers and 64-256 units in the dense layers. Six unit number combinations were generated by alternate doubling of one of those values at each step, starting with the number of convolutional units. We tested the following networks with sum merge, filter size of 15 and input dropout rates of 0 and 0.2-0.3 unless stated otherwise:

- *full*-RC, max pooling, 1 conv. and 1 dense layers
- *full*-RC, average pooling, 1 conv. and 1 dense layers
- *full*-RC, max pooling, 2 conv. and 2 dense layers
- *full*- and *siam*-RC, average pooling, 2 conv. and 2 dense layers

- *full*-RC, average pooling, 3 conv. and 3 dense layers, input dropout rate 0.2-0.25

We also applied batch normalization (Ioffe and Szegedy, 2015) to the RC-CNNs with 2 convolutional and 2 dense layers of 512 and 256 units (hereafter: 2x2-XL), and tested changing the filter size to 7 or 11 by training *full* RC-CNNs with the same layout. In addition, we trained two traditional CNNs without RC parameter sharing. One was a 2x2-XL architecture with the input dropput of 0.25, and the other was a 1x1 max pooling network with 256 convolutional and 128 dense units. A traditional LSTM was trained with 1 layer of 384 units and an input dropout rate of 0.2. Finally, we evaluated hybrid *full* and *Siamese* RC-networks with a convolutional layer of 512 units and filter size of 7, 11 or 15, followed by a recurrent layer of 384 units, trained with the input dropout of 0.2. Those networks were trained for just 10 epochs due to high computational cost.

#### 2.5 Loss function and training sets

We use binary cross-entropy for all training runs. We tested the effect of adding an $L_2$ regularization term while training both the *full* and *Siamese* RC-LSTMs with sum merge, 1 layer of 384 units and an input dropout rate of 0.2. We used regularization rates of $10^{-2}$, $10^{-3}$ and $10^{-5}$. Next, we tested using an imbalanced training set and weighting the errors by the inverse of the relative class frequency. To this end, we trained *full* and *Siamese* RC-LSTMs with sum merge, 1 layer of 384 units and input dropout rates between 0.1 and 0.3.

### 3 Results

#### 3.1 Reverse-complementarity constraint

We show predictions for each read in the test set against predictions for its reverse-complement in the Figure S1. Note that the reads are simulated from either of the strands at random. Even though the predicted pathogenic potentials are highly correlated (Spearman's $\rho > 0.999$, $p < 10^{-6}$) and identically distributed ($p > 0.999$, two-sample KS test) for both orientations, they are not equal. The differences are striking especially for the LSTM model. For the RC neural networks, the predictions are guaranteed to be exactly the same. Interestingly, the prediction distributions are very different for the CNN and LSTM models. We can only speculate that this could be related to the specific properties of the architectures – while the CNNs assume translational invariance and hence focus on localized sequence motifs, the LSTMs are able to also detect long-range relationships between more distant fragments of a read.

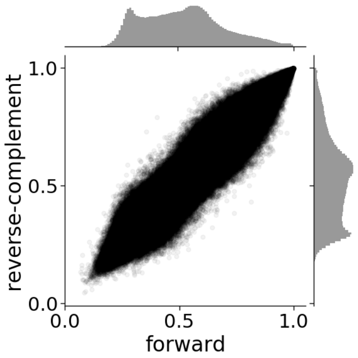


(a) CNN

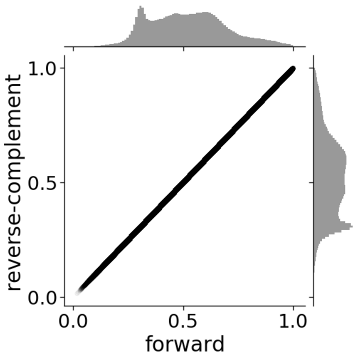


(b) RC-CNN

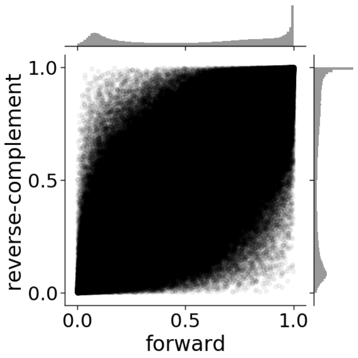


(c) LSTM

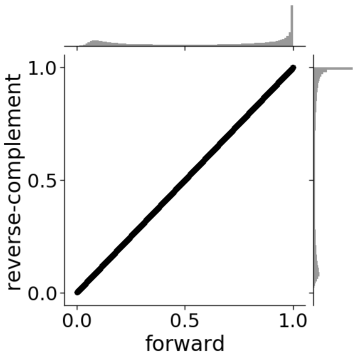


(d) RC-LSTM

Figure S1: Prediction distributions for traditional (S1a, S1c) and reverse-complement networks (S1b, S1d). Predictions for each of the reads (x-axis) are plotted against the predictions for reverse-complements of the same reads (y-axis). The RC-architectures guarantee identical predictions for both strands, which can be strikingly divergent for the traditional LSTM.

#### 3.2 Time performance

The prediction speed is difficult to compare, as PaPrBaG can only run on a CPU, and our neural networks are most efficiently used on GPUs. Both can be trivially parallelized depending on the number of devices available. We measured the time PaPrBaG needed for a complete prediction task (feature extraction and prediction itself) on a single Intel(R) Xeon(R) E7-4890 v2 CPU. The original version of PaPrBaG was able to classify 107 reads/s, and 47% of that time was used for feature extraction. The DNA-only version classified up to 167 reads/s, and most of the speed-up came from the reduction of feature extraction time (which was 25% of the total elapsed time). The classification performance was nearly identical.

The reverse-complement neural networks were tested on a single, consumer-level GPU (RTX 2080 Ti). We measured the time of prediction from sequences already converted to binary tensors, as the conversion step may be performed independently on separate CPU devices. The selected RC-LSTM model can predict up to 1817 reads/s and the RC-CNN reaches a speed of 4010 reads/s. Using this rapid architecture, a million-read sample may be processed in just over 4 minutes even on a desktop computer.
